# Microglia-targeted dendrimer-2PMPA therapy robustly inhibits GCPII and improves cognition in a mouse model of multiple sclerosis

**DOI:** 10.1101/2020.04.22.055228

**Authors:** Kristen Hollinger, Anjali Sharma, Carolyn Tallon, Lyndah Lovell, Ajit G. Thomas, Xiaolei Zhu, Siva P. Kambhampati, Kevin Liaw, Rishi Sharma, Camilo Rojas, Rana Rais, Sujatha Kannan, Rangaramanujam M. Kannan, Barbara S. Slusher

## Abstract

Roughly half of all individuals with multiple sclerosis (MS) experience cognitive impairment, but there are no approved treatments that target this aspect of the disease. Recent studies link reduced brain *N*-acetylaspartylglutamate (NAAG) levels to impaired cognition in various neurological diseases, including MS. NAAG levels are regulated by glutamate carboxypeptidase II (GCPII), which hydrolyzes the neuropeptide to *N*-acetyl-aspartate (NAA) and glutamate. Although several GCPII inhibitors, such as 2-(phosphonomethyl)-pentanedioic acid (2-PMPA), elevate brain NAAG levels and restore cognitive function in preclinical studies when given at high systemic doses or via direct brain injection, no GCPII inhibitors are clinically available due to poor bioavailability and limited brain penetration. Systemic hydroxyl dendrimers (~4 nm) have been successfully used to enhance brain delivery of drugs selectively to activated glia. We recently discovered that GCPII is highly upregulated in activated microglia after brain injury. To determine if dendrimer conjugation could enhance the brain delivery of GCPII inhibitors, specifically in the context of MS, we attached 2-PMPA to hydroxyl polyamidoamicne (PAMAM) dendrimers (D-2PMPA) using a highly efficient click chemistry approach. Targeted uptake of D-2PMPA into activated glia was subsequently confirmed in glial cultures where it showed robust anti-inflammatory activity, including an elevation in TGFβ and a reduction in TNFα. Given these positive effects, D-2PMPA (20mg/kg) or vehicle dendrimer were dosed twice weekly to experimental autoimmune encephalomyelitis (EAE)-immunized mice starting at disease onset (therapeutic paradigm). D-2PMPA significantly improved cognition in EAE as assessed by Barnes maze performance, even though physical severity was not impacted. Glial target engagement was confirmed, as CD11b+ enriched cells isolated from hippocampi in D-2PMPA-treated mice exhibited almost complete loss of GCPII activity. These data demonstrate the utility of hydroxyl dendrimers to enhance brain penetration and support the development of D-2PMPA to treat cognitive impairment in MS.

**Funding:** This work was funded by the National Multiple Sclerosis Society (RG-1507-05403 to BSS), the National Institute of Health NINDS (R01NS093416 to SK, RM and BSS), and Ashvattha Therapeutics. We would also like to acknowledge support for the statistical analysis from the National Center for Research Resources and NIH NCATS (1UL1TR001079).

**Highlights:**

- The GCPII inhibitor 2-PMPA was conjugated to hydroxyl PAMAM dendrimers (D-2PMPA)
- D-2PMPA targeted activated glia in culture and displayed anti-inflammatory activity
- When dosed systemically to EAE mice, D-2PMPA inhibited CD11b+ cell GCPII activity
- When dosed systemically to EAE mice, D-2PMPA improved cognitive function

## 1. Introduction

Multiple sclerosis (MS) is an autoimmune disease of the central nervous system (CNS). There are over 15 disease-modifying therapies approved by the FDA to treat the various subtypes of MS, but none target cognitive impairment that affects over half of all MS patients. This unmet treatment need is critical to explore, as MS-related learning and memory impairments have a measurable negative impact on quality of life, cost of living, and empolyability^1–4^. Targeted therapies may offer differentiated approaches to improve efficacy, therapeutic window, and reduce side effects.

The enzyme glutamate carboxypeptidase II (GCPII) metabolizes the neuropeptide *N*-acetylaspartyl glutamate (NAAG) into *N*-acetyl aspartate (NAA) and glutamate5. NAAG is one of the most abundant neuropeptides in the mammalian brain6 and regulates glutamate neurotransmission through its agonist activity at the metabotropic glutamate receptor 3 (mGluR3)^7^. Previous studies have demonstrated a significant correlation between brain NAAG levels and cognition in MS patients, with higher NAAG levels associated with better cognitive function^8^. Furthermore, in an animal model of MS, the potent (IC_50_=300 pM) and selective GCPII inhibitor 2-(phosphonomethyl)-pentanedioic acid (2-PMPA) was shown to enhance brain NAAG levels and improve cognitive function^8,9^. Despite this strong therapeutic data, existing GCPII inhibitors have poor bioavailability and brain penetration and therefore are unsuitable for clinical translation to MS patients.

Nanotechnology has shown promise in the development of smart drug delivery systems for targeted brain delivery of therapeutics with poor pharmacokinetic profiles^10^, with some undergoing translation^11^. Among the nanoparticles for biological applications, hydroxyl dendrimers have shown promise as targeted intracellular delivery systems, due to their size (~4-10 nm) and surface attributes^12^. Dendrimers are monodispersed and multivalent macromolecules with tailorable surface functionalities upon which bioactive molecules such as drugs, targeting ligands and/or imaging dyes can be covalently conjugated. Previous studies have demonstrated that systemically administered hydroxyl polyamidoamine (PAMAM) dendrimer-drug conjugates accumulate in activated glia at sites of brain inflammation, without a need for targeting moieties in multiple animal models including cerebral palsy, inflammatory preterm birth injury, and Rett syndrome^13–18^. These dendrimers take advantage of disease pathology to cross the impaired BBB and diffuse in the brain parenchyma, for increasingly phagocytic activated glia to take them up^14,16^. Moreover, the dendrimer-drug conjugates are consistently several-fold more potent and efficacious when directly compared to the equivalent amount of the free drug, likely due to their targeted delivery^19,20^. We therefore hypothesized that conjugating 2-PMPA to a dendrimer would enable preferential glial uptake in EAE-immunized mice, where GCPII is robustly upregulated in injured and diseased states^21^, making dendrimers a potentially efficacious GCPII inhibitor delivery vehicle for the treatment of cognitive impairment in MS.

Herein, we conjugated 2-PMPA to the surface of hydroxyl PAMAM dendrimers (D-2PMPA) using click chemistry and evaluated their glial uptake and anti-inflammatory efficacy in glial cultures and EAE-immunized mice.

## 2. Materials and methods

### 2.1 Synthesis of D-2PMPA and Cy5-D-2PMPA conjugates

Reagents and solvents were purchased from Sigma Aldrich or Fisher Scientific and were used as received unless otherwise stated. 2-PMPA (**compound 1**) was purchased from Sigma Aldrich (St. Louis, MO). Ethylenediamine-core PAMAM-OH dendrimer generation 4 having 64 hydroxyl end-groups, Pharma grade (**compound 3**) was received as a methanolic solution from Dendritech. Before use, the methanol was evaporated and dendrimer was further purified to remove generational impurities by dialysis. The dendrimer was solubilized in water and dialyzed against water using dialysis membrane of 3kDa. The dialysis membranes were purchased from Spectrum Laboratories Inc. Azido-PEG-11-alcohol was purchased from Broad Pharm and Cy5 NHS ester was purchased from GE healthcare and used as received. Deuterated solvents for NMR spectroscopy such as (DMSO‑*d6*), methanol (CD_3_OD), water (D_2_O) and chloroform (CDCl_3_) were purchased from Sigma. D-Cy5 was synthesized using our previously published protocol^18,22^.

#### Synthesis of compound 2

To a stirring solution of 2-PMPA (500mg, 2.211 mmoles) in anhydrous DMF (5mL), DMAP (300mg, 2.455 mmoles) and EDC (577, 3.00 mmoles) were added. The reaction mixture was stirred for 10 minutes. This was followed by the addition of azido-PEG-11-alcohol (1.06g, 2.010 mmoles). The solution was stirred at room temperature overnight. Upon completion, the DMF was evaporated and the crude product was purified using reverse-phase column purification using acetonitrile and water. The pure fractions were evaporated to afford compound **2** as viscous liquid. Yield: 41% **_1_H NMR** (500 MHz, D_2_O) δ 4.27 (t, 2H), 3.78 (t, 2H), 3.70 (s, 40H), 3.50 (t, 2H), 2.79-2.69 (m, 1H), 2.50 (t, 2H), 2.18 – 2.07 (m, 1H), 2.08-1.92 (qd, 2H), 1.92 – 1.79 (m, 1H). (**Supplementary Figure 1**) HRMS: m/z: calculated: 735.72, found: 758.31 [M+Na]+ (**Supplementary Figure 2**)

#### Synthesis of compound 4

To a stirring solution of D-OH (**3**, 2.00g, 0.140 mmoles) in anhydrous DMF, 5-hexynoic acid (235mg, 2.10 mmoles) followed by the DMAP (256mg, 2.10 mmoles) and EDC (537mg, 2.801 mmoles) were added. The stirring was continued for 24 hours at room temperature. The reaction mixture was then diluted with DMF and dialyzed against DMF for 6 hours followed by the overnight water dialysis. The solvents were changed frequently during dialysis. The aqueous solution was then lyophilized to obtain compound **4** as white solid. Yield: 91% **_1_H NMR** (500 MHz, D2O) δ 7.93 (m, D-internal amide *H*), 4.72 (bs, D-O*H*), 4.01 (t, ester –C*H*_2_), 3.49 – 3.21 (m, D- –C*H*_2_), 3.11 (m, D and linker–C*H*_2_), 2.79 – 2.58 (m, D and linker–C*H*_2_), 2.49-2.30 (m, D–C*H*_2_), 2.31-2.05 (m, D–C*H*_2_), 1.69 (t, linker–C*H*_2_). (**Supplementary Figure 3**)

#### Synthesis of compound 5

To a stirring solution of compound **4** (1.9g, 0.123mmoles) and Compound **2** (1.178g, 1.604 mmoles) in DMF:THF (1:1), CuSO_4_.5H_2_O (16.844mg, 0.067 mmoles) dissolved in 1mL water was added. This was followed by the addition of sodium ascorbate (26.804mg, 0.135 mmoles) dissolved in 1mL water. The reaction was carried out at 50°C in a microwave reactor for 6 hours. Upon completion, the reaction mixture was diluted with water, EDTA solution (1mL) was added and stirred for 1 hour. The solution was then transferred to a dialysis membrane (cut-off 1000kDa) and dialyzed against water for 15 hours. The aqueous solution was then lyophilized to afford the pure product as white solid. Yield: 82% **_1_H NMR** (500 MHz, D_2_O) δ 7.87 (s, triazole *H*), 4.58 (t, linker –C*H*_2_), 4.15 (t, linker – C*H*_2_), 3.92 (t, ester –C*H*_2_), 3.83 – 3.52 (m, PEG H and dendrimer –C*H*_2_), 3.53 – 3.23 (m, D- and linker–C*H*_2_), 3.16-2.65 (m, D- and linker–C*H*_2_), 2.66 – 2.34 (m, D–C*H*_2_), 2.24 – 2.06 (m, PMPA *H*), 2.04 – 1.84 (m, PMPA *H*), 1.85 – 1.54 (m, PMPA *H*). (**Supplementary Figure 4**) **HPLC:** Retention time: 22.99 minutes, purity: 99.69% (**Supplementary Figure 5**) **MALDI:** Theoretical 22,589; Found: 21,881 Da (**Supplementary Figure 6**)

#### Synthesis of compound 6

To a stirring solution of compound **4** (810mg, 0.053 mmoles) in DMF (10mL), GABA-BOC-OH (53.74mg, 0.265 mmoles) followed by DMAP (33mg, 0.270 mmoles) and EDC (33mg, 0.427 mmoles) were added. The reaction was stirred at room temperature for 24 hours. The solution was then transferred to a dialysis membrane (cut-off 1000kDa) and dialyzed against DMF followed by water for 15 hours. The aqueous solution was then lyophilized to afford the pure product as white solid. Yield: 86% **_1_H NMR** (500 MHz, DMSO) δ 8.09-7.68 (m, D-internal amide *H*), 6.83 (s, N*H* BOC), 4.71 (bs, D-O*H*), 4.07-3.95 (m, ester –C*H*_2_ of both linkers), 3.49 – 3.21 (m, D- and linker–C*H*_2_), 3.21 – 2.99 (m, D–C*H*_2_), 2.99 – 2.83 (D–C*H*_2_), 2.83 – 2.56 (D- and linker–C*H*_2_), 2.47 – 2.33 (D–C*H*_2_), 2.31 – 1.98 (m, D- and linker–C*H*_2_), 1.59 (t, linker–C*H*_2_), 1.37 (s, BOC *H*). (**Supplementary Figure 7**)

#### Synthesis of compound 7

To a solution of compound **6** (200mg) in DCM (7mL), trifluoroacetic acid (3mL) was added. The solution was vigorously stirred at room temp for 12 hours. DCM was then evaporated and TFA was removed by co-evaporation with methanol. This process was repeated several times. The trace solvents were removed using high vacuum to afford compound **7** as TFA salt in quantitative yield. **1_H_ NMR** (500 MHz, DMSO) δ 8.67-8.13 (m, D-internal amide *H*), 7.91 (s, -N*H*_2_), 4.07-3.95 (m, ester –C*H*_2_ of both linkers), 3.69 – 3.26 (m, D- and linker–C*H*_2_), 3.26 – 2.93 (m, D- and linker–C*H*_2_), 2.87 – 2.57 (m, D- and linker–C*H*_2_), 2.40 (m, D-C*H*_2_), 1.79 (t, linker–C*H*_2_). (**Supplementary Figure 8**)

#### Synthesis of compound 8

To a stirring solution of compound **7** (220mg, 0.014 mmoles) in DMF, PMPA-PEG-azide (144.06mg, 0.196 mmoles) followed by a catalytic amount of CuSO_4_.5H_2_O (1mg) in water (1mL) was added. The reaction was stirred for 5 minutes. This was then followed by the addition of sodium ascorbate (2mg) in water (1mL). The reaction was placed in the microwave at 50°C for 8 hours. Upon completion, the reaction mixture was diluted with water, and EDTA solution (1mL) was added and stirred for 1 hour. The solution was then transferred to a dialysis membrane (cut-off 1000kDa) and dialyzed against water for 15 hours. The aqueous solution was then lyophilized to afford the pure product as white solid. Yield: 79% **_1_H NMR**(500 MHz, DMSO) δ 8.30-7.74 (m, D-amide *H* and triazole *H*), 4.47 (t, linker–C*H*_2_), 4.06 (m, ester –C*H*_2_), 3.79 (t, linker –C*H*_2_), 3.58 – 3.28 (m, D- and linker– C*H*_2_), 3.24 – 2.98 (m, D- and linker–C*H*_2_), 2.93 – 2.59 (m, D- and linker–C*H*_2_), 2.40 – 1.98 (m, D–C*H*_2_), 1.96 – 1.38 (m, PMPA *H* and linker *H*). (**Supplementary Figure 9**)

#### Synthesis of compound 9

To a stirring solution of compound **8** (130mg, 0.005 mmoles) in DMF (5mL), DIPEA (0.1mL) followed by Cy5-NHS ester (5.06mg, 0.008 mmoles) dissolved in DMF (1mL) was added. The reaction was stirred at room temperature for 24 hours. The solution was then transferred to a dialysis membrane (cut-off 1000kDa) and dialyzed against DMF followed by water for 10 hours. The aqueous solution was then lyophilized to afford the pure product as blue solid. Yield: 86% **_1_H NMR** (500 MHz, DMSO) δ 8.41-8.70 (m, Cy5 *H*), 8.25 – 7.71 (m, D-amide *H* and triazole *H*), 7.77-7.60 (m, Cy5 *H*), 7.36-7.28 (m, Cy5 *H*), 6.62-6.51 (m, Cy5 *H*), 6.31-6.26 (m, Cy5 *H*), 4.47 (t, linker –C*H*_2_), 4.06 (m, ester –C*H*_2_), 3.79 (t, linker –C*H*_2_), 3.70 – 3.22 (m, D- and linker–C*H*_2_), 3.11– 2.59 (m, D- and linker–C*H*_2_), 2.32-1.97 (m, D–C*H*_2_), 1.92-1.75 (m, PMPA *H* and linker *H*), 1.75 – 1.35 (m, PMPA *H*). (**Supplementary Figure 10**) **HPLC:** Purity 99.9%, Retention time: 22.5 minutes (**Supplementary Figure 11**)

### 2.2 Compound characterization

#### Nuclear Magnetic Resonance (NMR) spectroscopy

_1_H NMR spectra were logged at 500MHz using Bruker spectrometer at 25°C. The chemical shifts of the residual protic solvent were reported in ppm relative to the internal trimethylsilane standard (δ = 0 ppm); CDCl_3_ (_1_H, δ = 7.27 ppm; 13C, δ = 77.0 ppm (central resonance of the triplet)), D_2_O (_1_H, δ = 4.79 ppm); and DMSO-*d*6 (1H, δ = 2.50 ppm) were used for chemical shifts calibration. The chemical shift multiplicities are abbreviated as follows: s = singlet, d = doublet, t = triplet, q = quartet, m = multiplet, and br = broad.

#### High Performance Liquid Chromatography (HPLC)

The purities of the intermediates and the final conjugates were analysed using HPLC (Waters Corporation, Milford, MA) equipped with a 2998 photodiode array detector, a 2475 multi λ fluorescence detector, a 1525 binary pump, and an in-line degasser AF. The HPLC was interfaced with Waters Empower software. A C18 symmetry 300, 5μm, 4.6×250mm column from Waters was used. The HPLC chromatograms were recorded at 210nm (dendrimer and 2-PMPA absorption), for D-2PMPA and at 650nm wavelength (Cy5 absorption) for Cy5-D-2PMPA. A gradient flow was used using a mobile phase consisting of buffer A: 0.1% TFA and 5% ACN in water and buffer B: 0.1% TFA in ACN). The gradient started from 100:0 (A:B) gradually increasing to 50:50 (A:B) at 20 min, finally returning to 100:0 (A:B) at 40 minutes maintaining a flow rate of 1 mL/min. For the purification of drug linker, semi-preparative HPLC from Shimadzu was used using same method with a flowrate of 5mL/min.

#### Mass spectroscopy

High resolution mass spectrometry (HRMS) was performed on Bruker microTOF-II mass spectrometer using ESI in the positive mode and direct flow sample introduction in CH_3_CN/ H_2_O (9:1) solvent system. The empirical formula confirmation was obtained by protonated molecular ions [M + nH]n+ or adducts [M + nX]n+ (X = Na).

Matrix assisted laser desorption ionization time of flight (MALDI-TOF) experiments were performed on Bruker Autoflex MALDI-TOF instrument using laser power of 55-100%. The sample was dissolved in ultra-pure water (4mg/mL) and the matrix was dissolved in acetonitrile:water mixture [50:50 (v/v)] at 10mg/mL concentration. 10μL of the dendrimer solution was mixed with 10μL of the matrix solution to prepare the samples, out of which 3 μL of the solution was spotted on a MALDI plate.

#### Dynamic light scattering (DLS) and Zeta potential (ζ)

The size and the zeta potential distribution of the D-2PMPA were analysed using Zetasizer Nano ZS (Malvern Instrument Ltd. Worchester, U.K.) equipped with a 50mW He-Ne laser (633 nm). The size measurements in triplicates were performed. The sample for size distribution measurement was prepared by dissolving the D-2PMPA in deionized water at a concentration of 0.5mg/mL. The solution was filtered through 0.2μm syringe filters (Pall Corporation, 0.2μm HT Tuffryn membrane). The measurement was performed in a UV transparent disposable cuvette having dimensions as 12.5 × 12.5 × 45mm (SARSTEDT). The zeta potential measurements were also performed in triplicates using a sample concentration of 0.2mg/mL in 10mM NaCl after the sample was filtered through 0.2μm syringe filters.

### 2.3 Primary glial cultures

Primary mixed glial cells were collected from neonatal rabbits (postnatal day 1) based on our previously established protocol^23^. Mixed glia were maintained in DMEM media containing 4.5 g/L glucose and 1.4 mM L-glutamine (Corning Cellgro, Manassas, VA USA) with 10% FBS and 1% pen/strep antibiotic at 37°C and 5% CO_2_ atmosphere.

### 2.4 Uptake studies in primary glial cultures

*In vitro* uptake of fluorescently labeled Cy5-D-2PMPA was assessed in primary mixed glial cultures from neonatal rabbit brain as previously reported^23^. Cells were plated into glass-bottom culture dishes and treated with 50μg/mL Cy5-D-2PMPA in 5% serum medium. At particular time points (3, 6, 12, 18 and 24 hours), the cells were fixed with 2% PFA for 15 minutes. The fixed cells were blocked with 5% normal goat serum for 4 hours followed by incubation with anti-Iba-1 antibody (1:300, Abcam, Cambridge, UK) for 12 hours at 4°C to label microglial cells. Next the cells were washed with tris buffer with 0.1% triton X and were incubated with secondary antibody goat anti mouse Cy3 (1:500, Invitrogen, Rockland, IL, USA) for Iba-1 and anti-GFAP (1:500, eBioscience, San Diego, CA, USA) overnight at 4°C to stain astrocytes. The cells were washed twice with PBS for 5 minutes, stained with 4’,6-diamidino-2-phenylindole (DAPI) (1:1000) (Invitrogen, Grand Island, NY, USA) for 15 minutes, and imaged under an LSM 710 confocal microscope (Carl Zeiss, Hertfordshire, UK) for identification of the dendrimers (Cy5-D-2PMPA) in microglia and astrocytes.

### 2.5 Anti-inflammatory effects in LPS-treat primary glial cultures

Primary rabbit mixed glial cells were seeded onto 12-well plates. After 48 hours, cells were activated with lipopolysaccharides (LPS; lot# 127M4130V, Sigma, St. Louis, MO USA) at 300 EU/mL for 6 hours. After 6 hours, the cells were treated with various concentrations of D-2PMPA for 24 hours. For viability analyses, 5mg/mL MTT solution was added for 4 hours and samples were analyzed at 540nm. For rt-qPCR, cells were incubated in fresh media for 24 hours, followed by collection into Trizol (Invitrogen, Carslbad, CA USA) and RNA extracted per manufacturer’s instructions. RNA was then converted into cDNA for rt-qPCR analysis. Primer sequences:

TGFβ: F: TGAGAGGTGGAGAGGAAATAGA, R: GGAACTGATCCCGTTGATGT
mGluR3: F: CGACAAGTCTCGCTACGATTAC, R: CACGTAGGTCCAGTTGAAGAAG
NR2A: F: CAAGGATCCCACGTCTACTTTC, R: AAGACGTGCCAGTCGTAATC
TNFα: F: TAGTAGCAAACCCGCAAGTG, R: CTGAAGAGAACCTGGGAGTAGA
iNOS: F: CAGGACCACACCCCCTCGGA, R: AGCCACATCCCGAGCCATGC
GAPDH: F: TGACGACATCAAGAAGGTGGTG, R:
GAAGGTGGAGGAGTGGGTGTC

### 2.6 Mice

Female 7-week-old C57BL/6J mice were purchased from Jackson Laboratory (Bar Harbor, ME) and housed in the Miller Research Building Johns Hopkins animal facility. All protocols were approved by the Johns Hopkins Institutional Animal Care and Use Committee and cared for in compliance with the National Institutes of Health guide for the care and use of Laboratory animals (NIH Publications No. 8023, revised 1978)

### 2.7 EAE immunizations and scoring

8-10-week-old mice were immunized for EAE as previously described8 with minor modifications. Briefly, mice were administered murine MOG 35-55 (Johns Hopkins Peptide Synthesis Core Facility, Baltimore) in incomplete Freund’s adjuvant (Sigma-Aldrich) supplemented with heat-killed mycobacterium tuberculosis (Sigma-Aldrich) via two subcutaneous flank injections. On days 0 and 2, mice received an intraperitoneal injection of 250 ng pertussis toxin (List Biological Laboratories). EAE disease scores were assigned by an observer blinded to the treatment as previously described^9^ based on the following scale with 0.5 increments for intermediate scores: 0=normal, 1=limp tail, 2=wobbly gait, 3=dragging hind flank, 4=hind limb paralysis, 5=quadriplegia. Experiments were conducted in duplicate.

### 2.8 D-2PMPA administration

To evaluate the effects of D-2PMPA treatment, vehicle (empty dendrimer) and D-2PMPA solutions in 0.9% sterile saline were prepared fresh every 2 weeks and stored at 4°C. Twice per week, mice (n=10) received intraperitoneal injections of vehicle or 20mg/kg D-2PMPA at a volume of 10μL/g body weight. Prior to treatment administration, solutions were brought to room temperature and vortexed.

### 2.9 Glial uptake studies in EAE-immunized mice

To evaluate *in vivo* dendrimer uptake, a separate cohort of 3 mice were immunized for EAE, sacrificed, and brains were processed and stained for immunohistochemical analysis. Mice were injected intraperitoneally with 55mg/kg Cy5-D-2PMPA conjugates on Day 14 post-immunization, and sacrificed 24 hours later via cardiac perfusion with ice cold PBS under anesthesia. Brains were dissected and post-fixed in 10% neutral buffered formalin for 48 hours at 4°C. Brains were then moved through 15% then 30% sucrose solutions (Thermo Fisher Scientific, Waltham, MA) for 48 hours at 4°C, frozen on dry ice, then stored at −80°C until sectioning. Brain tissues were sectioned using a cryostat (Microm HM 505E, International Medical Equipment, MI, USA) at a thickness of 30 μm. Sections were permeabilized and blocked for 1h at room temperature in 0.3% Triton (Sigma-Aldrich)/5% goat serum (Jackson Laboratories), then stained overnight at 4°C with a primary antibody against microglia (Iba1; Wako, Richmond VA). Sections were then stained with the appropriate secondary antibodies for 1h at room temperature. The slides were then washed in 1X PBS 3 times for 5 minutes before being incubated with Hoechst 33342 (Thermo Fisher) for 5 minutes before being washed again and coverslipped with prolong diamond (Life Technologies). The slides were then imaged at 20x with a LSM800 confocal microscope (Zeiss). Images were processed using Zen Blue software (Zeiss).

### 2.10 Cognition Studies in EAE-immunized mice

To evaluate cognition in EAE mice, the Barnes maze test was administered as previously described_8_ approximately 5 weeks post-immunization. Briefly, mice were placed in the center of the maze (Maze Engineers, Glenview, IL) and trained to find a hidden target platform for 2 trials per day over 4 consecutive days. Primary latency, or the time lapsed between the start of the test and the mouse locating the target platform, and paths taken by the mice were automatically recorded. The researcher conducting cognition studies was blinded to the treatment groups.

### 2.11 Hippocampal CD11b+ cell isolations

Mice completing the Barnes maze cognitive test were delivered a terminal dose of dendrimer vehicle or D-2PMPA then sacrificed 24h later via cardiac perfusion with ice cold PBS. Hippocampi, defined by landmarks and neuroanatomical nomenclature in the atlas of Franklin and Paxinos (anteroposterior: −0.95 - −4.03 mm, mediolateral: ±3.75 mm from bregma, dorsoventral: −1.75 - −5.1 mm from the dura)^24^, were rapidly and bilaterally dissected on an ice-cold plate. CD11b+ cells were isolated from hippocampi as previously described^25^. Briefly, brain tissue was minced in HBSS (Sigma-Aldrich, St. Louis, MO) and dissociated with neural tissue dissociation kits (MACS Militenyi Biotec, Auburn, CA). After passing through a 70 μm cell strainer, homogenates were centrifuged at 300 g for 10 minutes. Supernatants were removed, cell pellets were resuspended, and myelin was removed by Myelin Removal Beads II (MACS Militenyi Biotec). Myelin-removed cell pellets were resuspended and incubated with CD11b MicroBeads (MACS Militenyi Biotec) for 15 minutes, loaded on LS columns and separated on a quadroMACS magnet. Cells were flushed out from the LS columns, then washed and resuspended in sterile HBSS (Sigma-Aldrich). Viable cells were counted with a hemacytometer and 0.1% trypan blue staining. Each brain extraction yielded approximately 5×10_5_ viable CD11b+ cells. Samples were stored at −80°C until GCPII activity assays were performed.

### 2.12 GCPII activity assay

GCPII activity measurements were carried out on isolated CD11b+ cells based on previously published methods^26,27^. To determine the IC_50_ values of 2-PMPA, 2-PMPA-PEG-azide, D-2PMPA and the unconjugated generation 4 dendrimer (G4-OH), reactions were carried out in the presence or absence inhibitors, NAA-[_3_H]-G (30 nM, 48.6 – 49.6 Ci/mmol) and human recombinant GCPII enzyme (40 pM final) in Tris-HCl (pH 7.4, 40 mM) and 1 mM CoCl_2_. The reactions were carried out at 37°C for 20 minutes and stopped with ice-cold sodium phosphate buffer containing 1mM EDTA (pH 7.4, 0.1 M, 50 μL). 90 μL aliquots from each terminated reaction was then transferred to 96-well spin columns containing AG1X8 ion-exchange resin and the plate centrifuged at 990 rpm for 5 minutes using a Beckman GS-6R centrifuge equipped with a PTS-2000 rotor. NAA-[_3_H]-G was bound to the resin and [_3_H]-G eluted in the flow through. To ensure complete elution of [_3_H]-G, columns were washed twice with formate (1 M, 90 μL). The flow through and the washes were collected and 200 μL aliquots transferred to a solid scintillator-coated 96-well plate (Packard) and dried to completion. The radioactivity corresponding to [_3_H]-G was determined with a scintillation counter (Topcount NXT, Packard, counting efficiency 80%). Subsequently, IC_50_ curves were generated from CPM results, using both Microsoft Office Excel 2016 and IDBS’s XLfit 5.5.0.5 macro embedded within Excel.

To confirm target engagement, glial cell pellets were suspended in 150μl ice-cold Tris buffer (40 mM, pH 7.5) containing protease inhibitors (Roche, Complete Protease Inhibitor Cocktail, 1 tablet in 5 ml) and sonicated using Kontes’ Micro Ultrasonic Cell Disrupter (three pulses of 10s duration on ice, 30s between pulses). The resulting homogenates were spun down (16,000 × g for 2 min at 4°C) and the supernatants collected for both GCPII activity and total protein analysis. GCPII reaction was initiated after the addition of cobalt chloride (1 mM) and NAA-[_3_H]-G (40nM) pre-warmed to 37°C. The reactions were carried out in 50 μl reaction volumes in 96-well microplates for 2h min at 37°C. At the end of the reaction period, the assay was terminated, un-hydrolyzed NAA-[_3_H]-G and [_3_H]-G separated, and radioactivity corresponding to [_3_H]-G measured as detailed above. Finally, total protein measurements were carried as per manufacturer’s instructions using BioRad’s Detergent Compatible Protein Assay kit and data presented as fmol/mg/h.

### 2.13 Statistical Analyses

Statistical analyses were completed using GraphPad Prism 6.0. One-way ANOVA with Tukey’s multiple comparisons post-hoc test measured differences in PCR studies. Repeated measures two-way ANOVA with Sidak’s multiple comparisons post-hoc test measured D-2PMPA treatment effects on EAE disease scores and Barnes maze cognitive performance. Unpaired t-tests determined statistical significance of GCPII activity in microglia. P values <0.05 were considered statistically significant. Statistical analyses assistance and consultation were provided by the Johns Hopkins Institute for Clinical and Translational Research Biostatistics Program.

## 3. Results

### 3.1 Synthesis and characterization of D-2PMPA conjugate

The D-2PMPA conjugate was prepared by the covalent attachment of 2-PMPA molecules on the surface of hydroxyl PAMAM dendrimers (D-OH) using copper (I) catalyzed click chemistry (CuAAC). The conjugation of 2-PMPA on the surface of dendrimers was achieved in three steps: (i) the modification of the drug to attach a linker; (ii) partial modification of the surface of dendrimer with another linker to bring complementary functional group and finally, (iii) clicking the drug-linker to the dendrimer linker. The linkers were attached both to the dendrimer and the drug; (a) to bring environment sensitive linkages for intracellular drug release; (b) to obtain complementary functional groups for conjugation and (c) to reduce the steric hindrance for optimal drug loading.

The synthesis of D-2PMPA began with the modification of 2-PMPA to attach a short polyethylene glycol (PEG) linker with azide focal point to participate in the click reaction (**Figure 1A**). 2-PMPA (**1**) was reacted with azido-PEG-11-alcohol using EDC, DMAP as coupling agents. Compound **2** was purified by the reverse phase column chromatography. On the other hand, the surface of D-OH (**3**) was partially modified by the attachment of alkyne linker (**Figure 1B**). D-OH was reacted with hexynoic acid to bring approximately 11 alkyne moieties on the surface of dendrimer **4**. The dendrimer surface was only partially modified to maintain the inherent targeting potential of the hydroxyl dendrimer. The number of attached linkers was calculated by comparing the integration of dendrimer internal amide protons to the newly formed ester methylene protons in _1_H NMR (**Figure 1C**). In the final step, D-hexyne (**4**) and 2-PMPA-PEG-azide (**2**) were clicked together via CuAAC click reaction using a catalytic amount of copper sulfate and sodium ascorbate to afford D-2PMPA (**5**). The confirmation of the click reaction was achieved by comparing proton NMR spectra of D-hexyne and PMPA-PEG-azide to D-2PMPA. The proton NMR spectra of D-2PMPA clearly shows the presence of 2-PMPA protons and shifts in the methylene protons adjacent to the triazole ring. The number of 2-PMPA molecules conjugated on dendrimer were analyzed by comparing 2-PMPA protons to dendrimer protons confirming the attachment of an average of 11 drug molecules with a loading of ~10% weight by weight. Due to the overlap with dendrimer internal protons, the appearance of triazole ring protons was not evident when NMR was taken in DMSO-*d6*. The _1_H NMR spectra of D-2PMPA clearly shows the appearance of triazole H (δ 7.87 ppm) in the spectra obtained in D_2_O confirming the success of conjugation. D-2PMPA was further analyzed by HPLC which showed a clear shift in retention time for the conjugate (23.08 minutes) from the starting alkyne (22.0 minutes) and azide (20.74 minutes, **Figure 1D**). The HPLC purity of D-2PMPA was >99.5% (**Supplementary Figure 5**). The size of D-2PMPA was 4.7 nm and zeta potential was nearly neutral (−2.07 mV) as analyzed by dynamic light scattering (**Figure 1E, F**).

**Figure 1:**
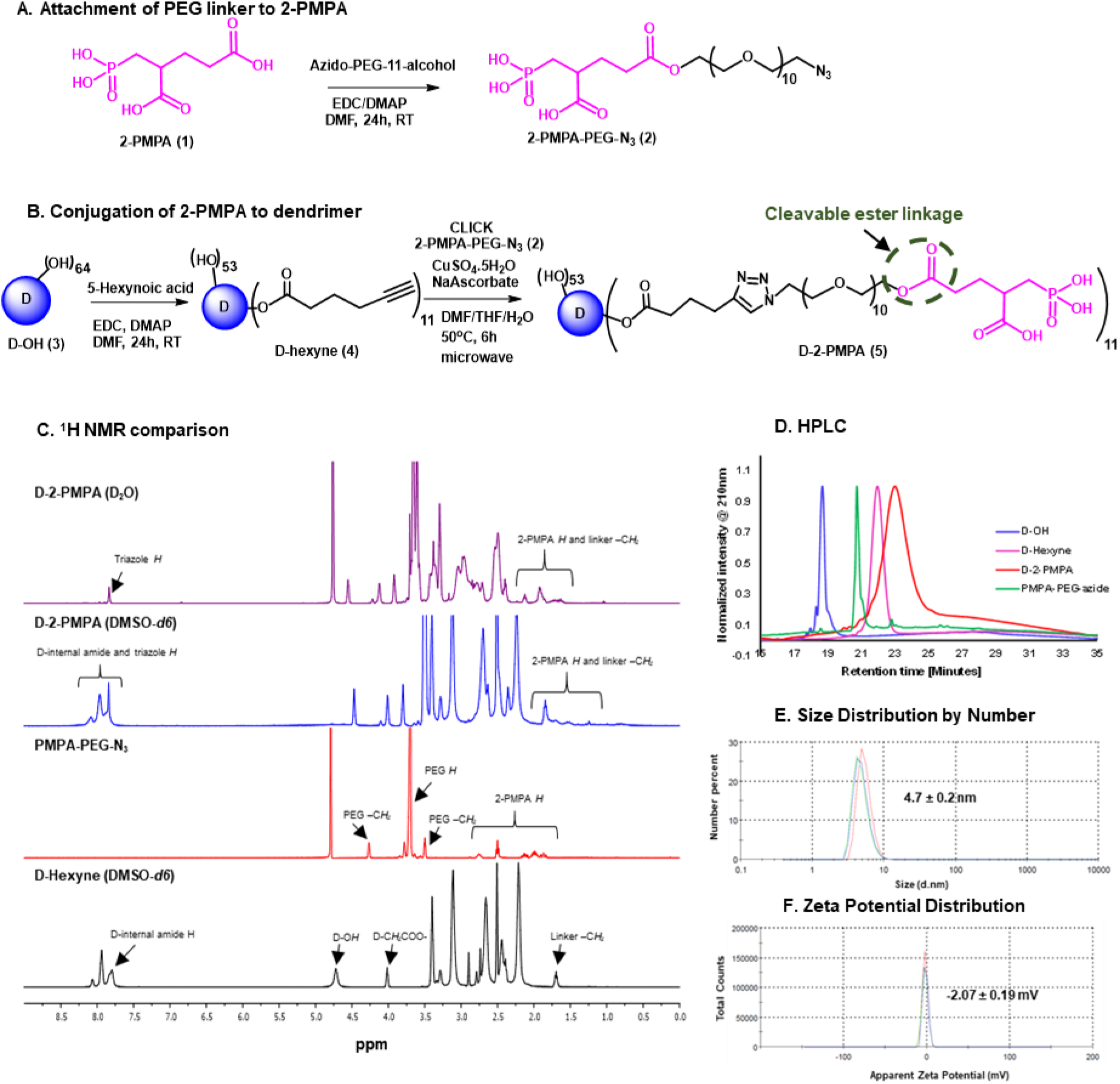
Synthesis and characterization of D-2PMPA. **A.** Synthetic protocol for the attachment of cleavable linker to 2-PMPA; **B.** The synthetic scheme for the conjugation of 2-PMPA on the surface of hydroxyl PAMAM dendrimer; **C.** Comparison of _1_H NMR spectra for the intermediates and the final D-2PMPA conjugate; **D.** HPLC chromatogram showing distinct shifts at each reaction step; **E.** Size distribution by number of D-2PMPA conjugate, and **F.** Zeta potential distribution of D-2PMPA as analyzed by the dynamic light scattering.

### 3.2 Synthesis and characterization of fluorescently-labeled Cy5-D-2PMPA conjugate

We further constructed fluorescently-labeled Cy5-D-2PMPA conjugates to perform *in vitro* and *in vivo* imaging to confirm that the attachment of 2-PMPA on the surface of dendrimer did not alter its targeting capabilities. The synthesis started with the modification of compound **4** to synthesize a multifunctional dendrimer with an amine group in addition to alkyne and hydroxyl groups (**Figure 2A**). This was achieved by the reaction of compound **4** with GABA-BOC-OH to obtain compound **6** with BOC protected amine which was then de-protected under mild acidic conditions using trifluoroacetic acid to yield multifunctional dendrimer **7**. The choice of the functional groups was made so that they did not interfere with each other during reactions. The alkynes in dendrimer **7** were further reacted with 2-PMPA-PEG-azide using classical click conditions as described earlier and the resulting compound **8** was obtained with approximately 11 PMPA molecules and an amine group. The dendrimer **8** was finally reacted with Cy5-NHS ester using activated acid amine coupling reaction to get Cy5-D-2PMPA. The structure of the final fluorescently-labeled conjugate was characterized using _1_H NMR spectroscopy (**Supplementary Figure 10**). The purity of Cy5-D-2PMPA was >99% as analyzed by HPLC (**Figure 2B** **and Supplementary Figure 11**). The UV/Vis spectra of Cy5-D-2PMPA showed the absorption at both dendrimer and 2-PMPA absorption wavelength (200nm) and Cy5 wavelength (650nm) further confirming the formation of the conjugate (**Figure 2C**).

**Figure 2:**
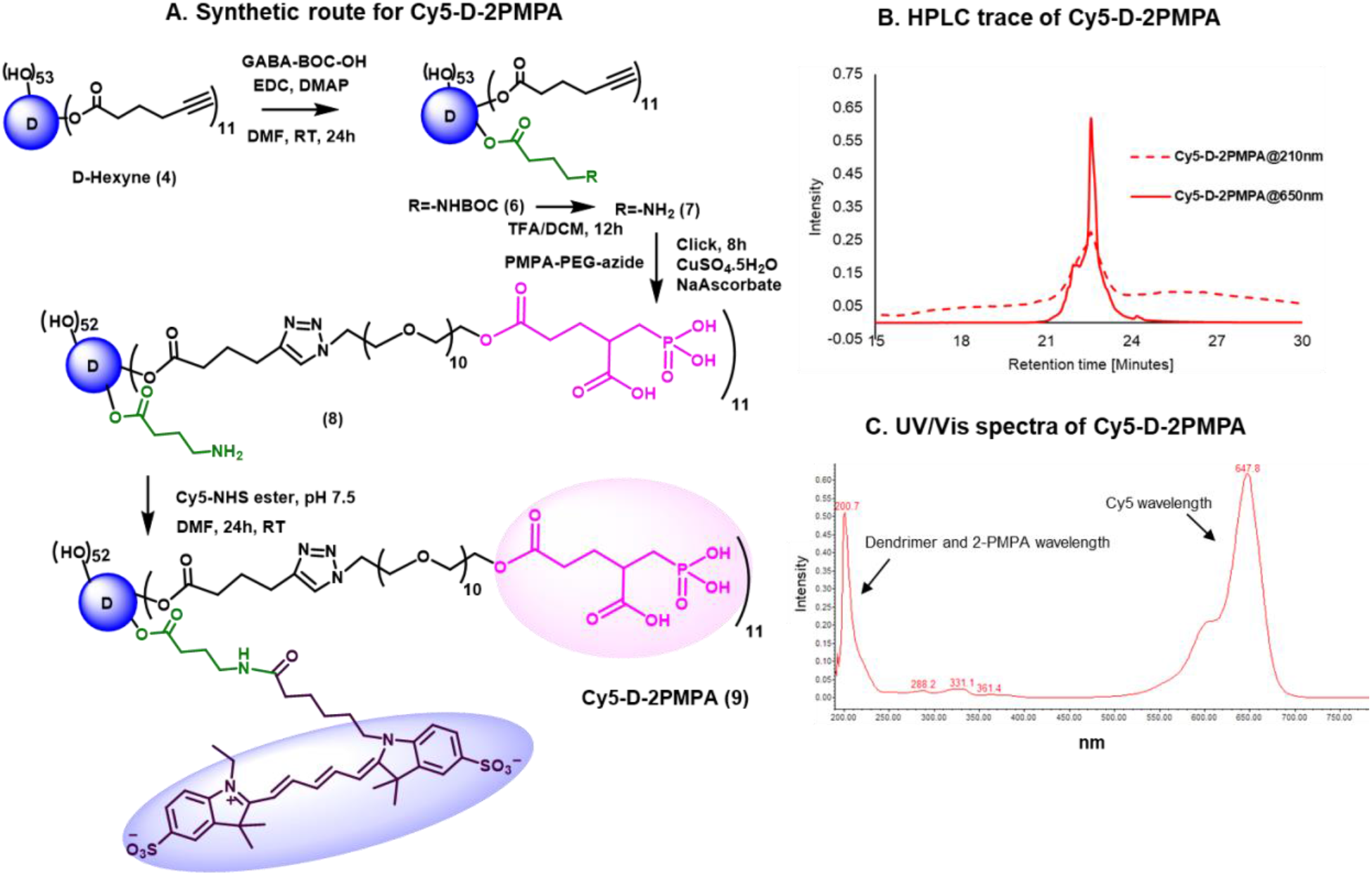
Synthesis and characterization of Cy5-D-2PMPA. **A.** Synthetic protocol for the synthesis of fluorescently-labeled dendrimer-2-PMPA conjugate (Cy5-D-2PMPA); **B.** HPLC chromatogram of Cy5-D-2PMPA at dendrimer absorption wavelength (210nm) and Cy5 absorption wavelength (650nm); and **C.** UV/Vis profile of Cy5-D-2PMPA showing absorption at both dendrimer and Cy5 wavelengths.

### 3.3 In vitro evaluation of D-2PMPA

#### 3.3.01 D-2PMPA inhibits human recombinant GCPII enzyme activity in vitro

To evaluate whether 2-PMPA conjugated to the dendrimer retained activity as an inhibitor of GCPII, inhibition was measured in the presence of D-2PMPA and compared with free 2-PMPA, 2-PMPA-PEG-azide, and D-OH (**Figure 3**). The IC_50_ of D-2PMPA was 3.50 ± 0.05nM, an approximate 10-fold loss of potency relative to free 2-PMPA, 0.20 ± 0.03nM. 2-PMPA conjugated to PEG-azide showed similar activity with an IC_50_ of 1.60nM ± 0.04nM, while negative control D-OH displayed no inhibitory activity.

**Figure 3:**
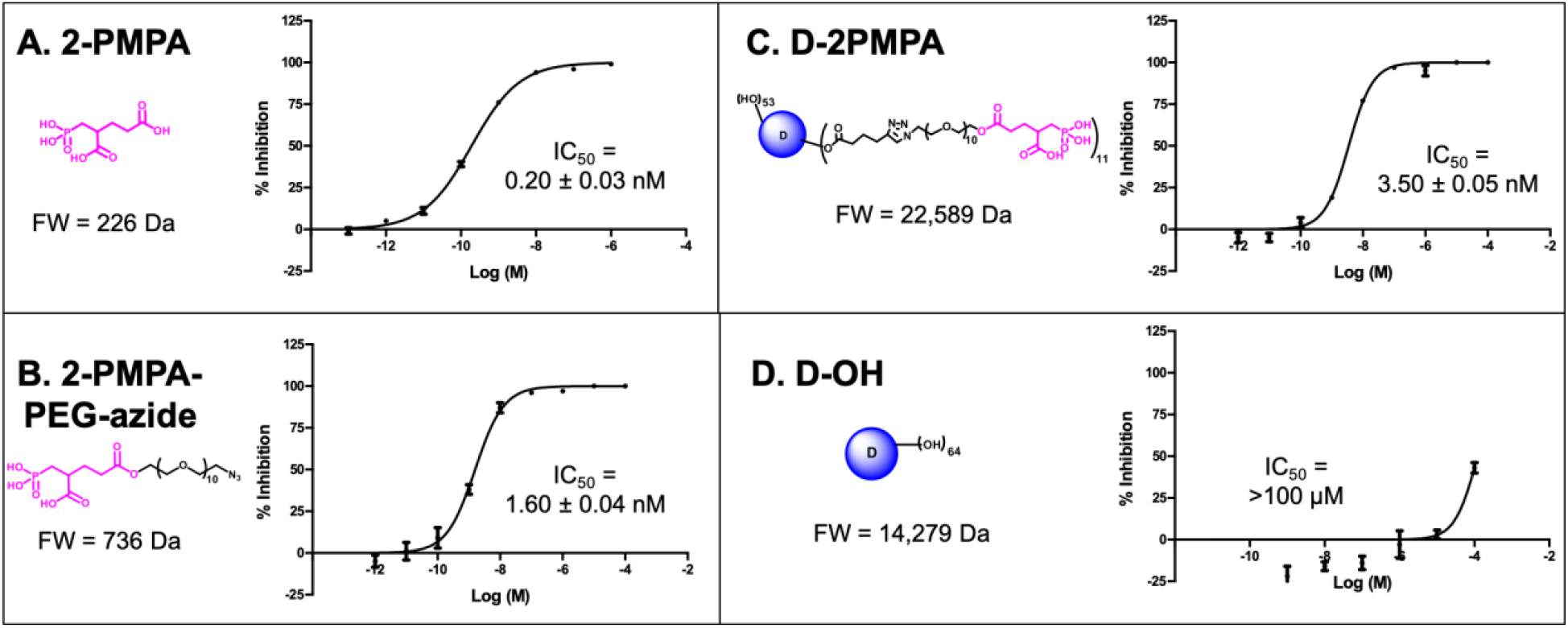
The comparative IC_50_s of the free drug, drug linker and dendrimers determined using human recombinant GCPII. **A.**2-PMPA; **B.** 2-PMPA-PEG-azide; **C.** D-2PMPA; and **D.** D-OH

#### 3.3.02 D-2PMPA localizes primarily to microglial cells in primary mixed glial cultures

We used mixed glial cultures containing both microglia and astrocytes to demonstrate the uptake kinetics of Cy5-D-2PMPA in both cell populations (**Figure 4**). At early time points (3 and 6 hours), Cy5-D-2PMPA were found predominantly co-localized by microglia (Iba-1+) cell population in the mixed glial culture (**Figure 4A-D** and **E-H**, white arrows). At 3 hours, ~50% of the Iba1+ microglial cells demonstrated dendrimer uptake (qualitatively), and at 6 hours, almost all the microglia population demonstrated dendrimer co-localization. This preferential uptake of Cy5-D-2PMPA by microglial population in mixed glial cells was similar to D-Cy5 (without conjugated PMPA) uptake as reported previously^23^ suggesting that conjugating 2-PMPA does not alter the cellular uptake of dendrimers. Twelve hours post Cy5-D-2PMPA treatment, microglial uptake was still evident, with a few of the GFAP+ astrocytes also demonstrating Cy5-D-2PMPA uptake (**Figure 4I-L**, white arrowheads respectively).

**Figure 4:**
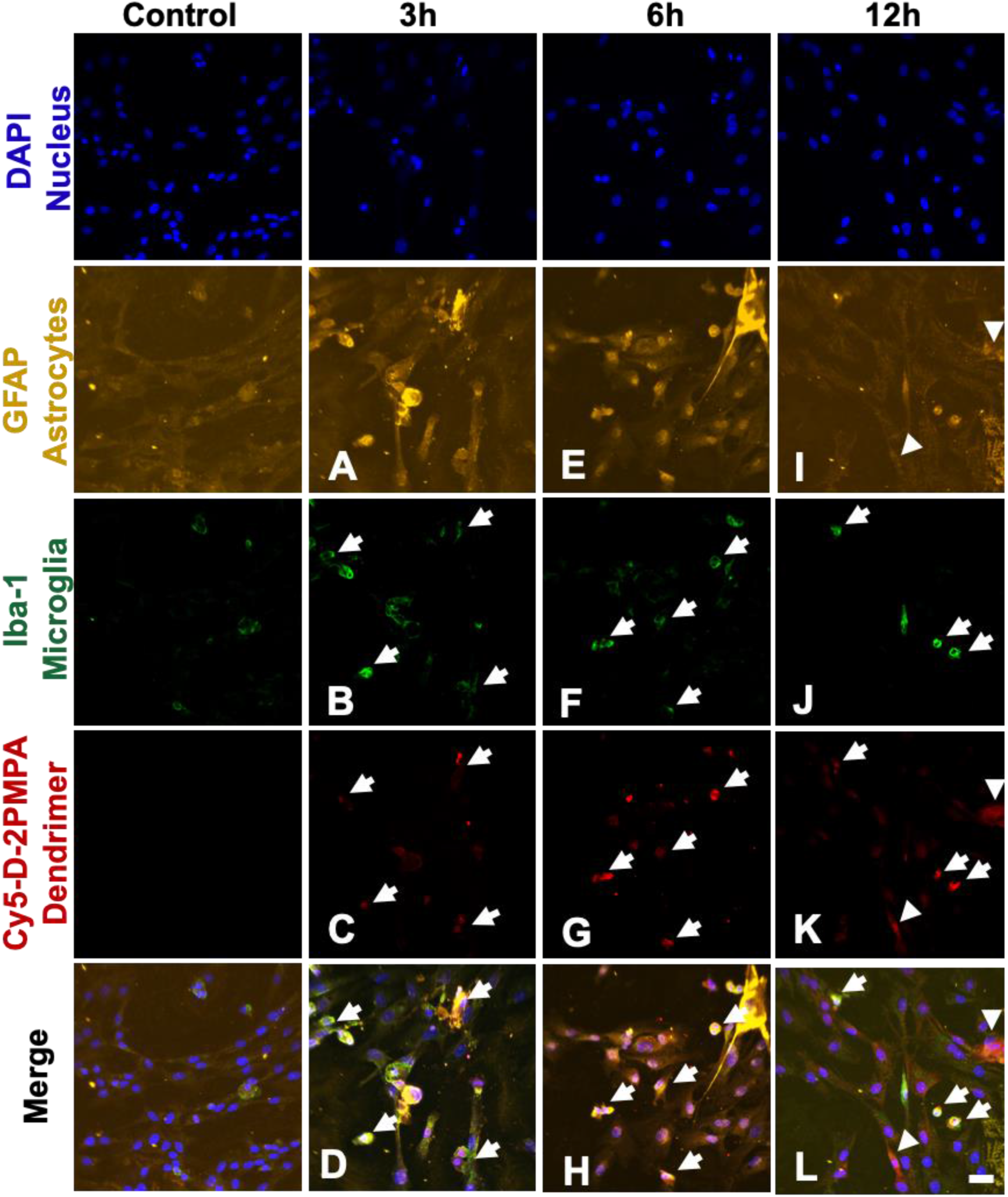
Confocal images of mixed primary glial culture demonstrating Cy5-D-2PMPA uptake. Microglia were stained using Iba-1 (green), astrocytes were stained using GFAP (orange), nuclei were stained using DAPI (blue), and the D-2PMPA conjugates were labelled with Cy5 (red). Cells were treated with 50 μg/mL of Cy5-D-2PMPA to evaluate uptake. At 3-6 hours post treatment, Cy5-D-2PMPA were preferentially taken-up by microglial cells (**A-D** and **E-H**, white arrows). Microglial uptake of Cy5-D-2PMPA continued at 12-hour post-treatment (**I-L**, white arrows), and astrocytes also demonstrated some signs of uptake (**I-L**, white arrowheads). Scale bar 20μm.

#### 3.3.03 D-2PMPA is anti-inflammatory in LPS-treated glial cultures

Mixed glial cultures were used to evaluate the *in vitro* activity of D-2PMPA. Dendrimer delivery of 2PMPA to LPS-treated glial cultures significantly upregulated TGFβ and mGluR3 (**Figure 5A-B**, P<0.05), and causes a trend towards an increase in NR2A (**Figure 5C**). D-2PMPA also lowered levels of the oxidative stress marker iNOS (**Figure 5D**) and significantly lowered levels of the pro-inflammatory cytokine TNFα (**Figure 5E**, P<0.05). MTT toxicity assay revealed no toxicities at the concentration used in these studies (**Figure 5F**).

**Figure 5:**
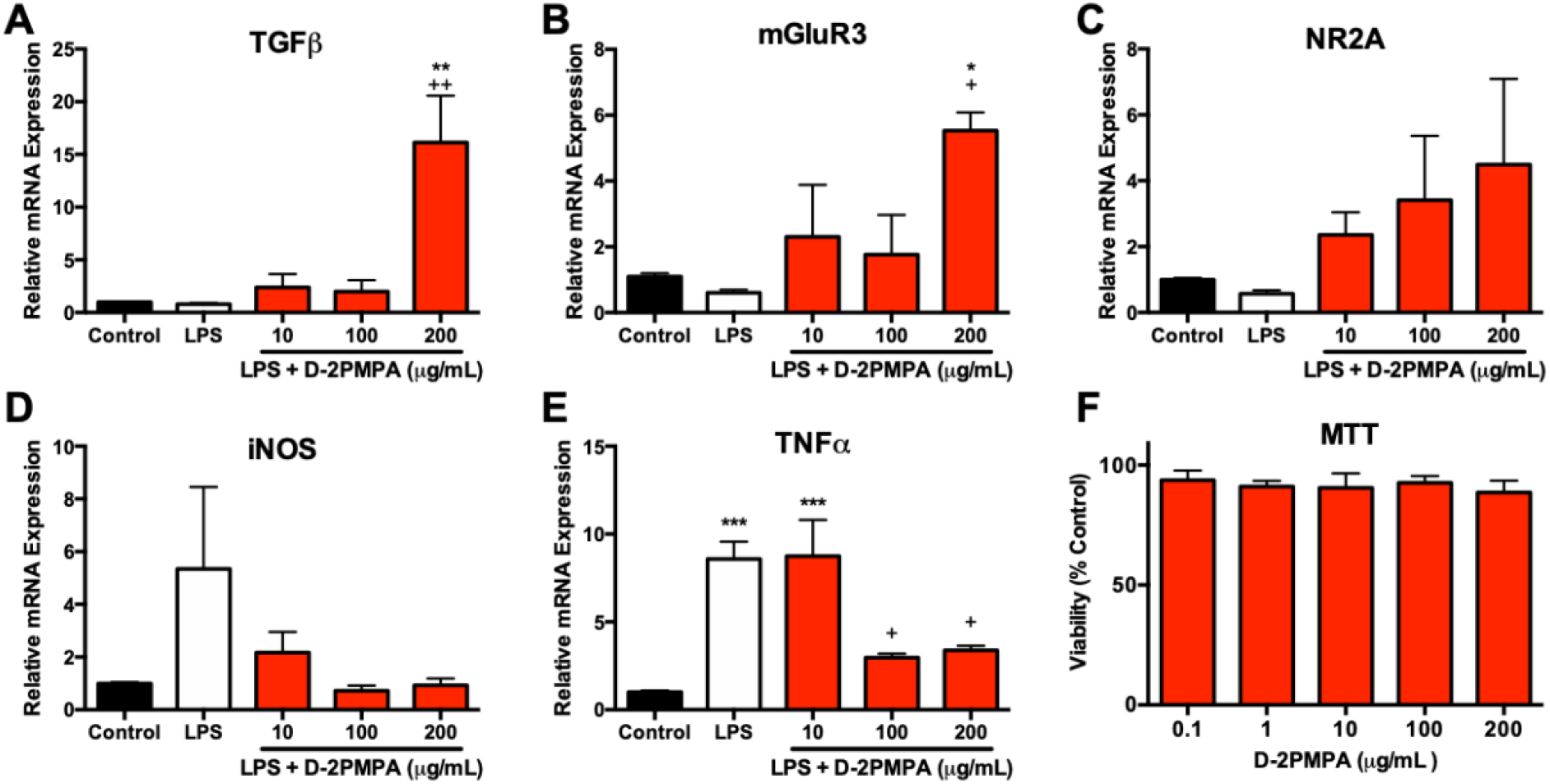
Effect of D-2PMPA on LPS-treated glial cultures. D-2PMPA upregulates TGFβ pathway mRNA, including **A.** TGFβ; **B.** mGluR3; and **C.** NR2A, and downregulates pro-inflammatory markers; **D.** iNOS; **E.** TNFα. **F.** D-2PMPA has no effect on cell viability as assessed by the MTT assay. Significantly different from control at P<0.05(*), P<0.01(**), P<0.001(***). Significantly different from LPS at P<0.05(+), P<0.01(++).

### 3.4 In vivo evaluation of D-2PMPA

#### 3.4.01 D-2PMPA is taken up by microglia in vivo

To evaluate D-2PMPA brain uptake following IP administration, EAE mice were administered a single IP Cy5-D-2PMPA dose on Day 14, sacrificed 24 hours later, and brains were imaged (**Figure 6**). Sections stained with Iba1 (red) confirmed perinuclear Cy5-D-2PMPA uptake in activated microglial cells. The highest concentrations of these positive cells were located in the molecular layer of the dentate gyrus near the third ventricle. No appreciable uptake was observed in astrocytes.

**Figure 6:**
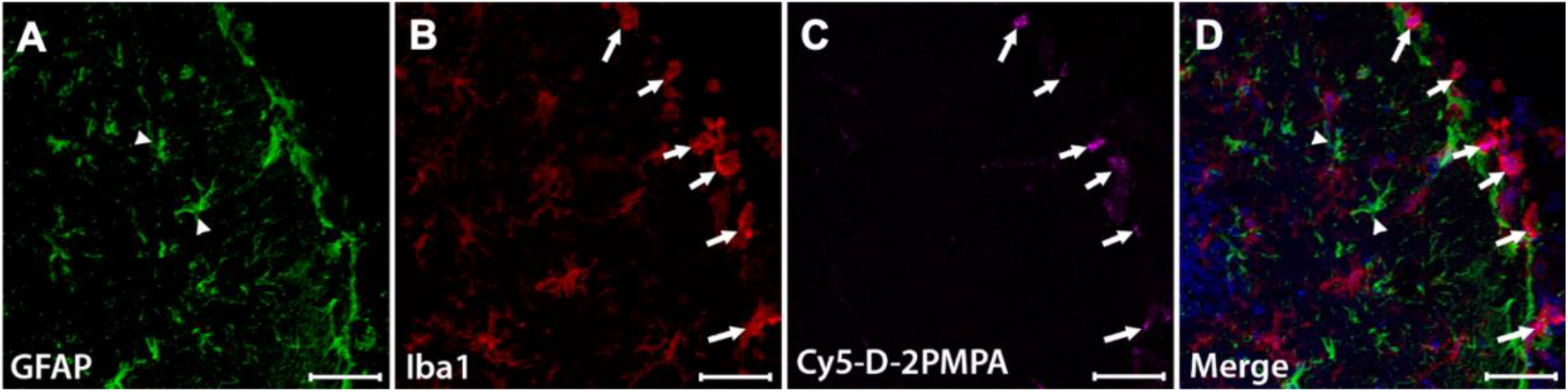
Cy5-D-2PMPA is engulfed by activated microglia. EAE mice were administered a single dose of D-2PMPA 14 days post-immunization and sacrificed 24 hours later. Representative brain images illustrate Iba1 positive activated microglia (**B**, red) engulfing Cy5-D-2PMPA (**C**, violet) (arrows) along the edge of the dentate gyrus, while GFAP positive astrocytes (**A**, green) have no Cy5-D-2PMPA signal (arrowheads). Nuclei are stained blue in the merged image (**D**). Scale bars, 50μm.

#### 3.4.02 D-2PMPA administration does not alter physical severity in EAE mice, but selectively improves cognitive function

To determine the effects of GCPII inhibition on physical severity of EAE, mice were administered biweekly injections of either D-2PMPA or empty dendrimer vehicle from the time of physical EAE disease onset. Mice developed signs of EAE approximately 2 weeks post-immunization. Data from two independent experiments demonstrate that inhibition of GCPII via D-2PMPA treatment does not affect the severity or progression of EAE, as indicated by no detectable differences in EAE scores throughout the duration of the experiment (**Figure 7A**).

**Figure 7:**
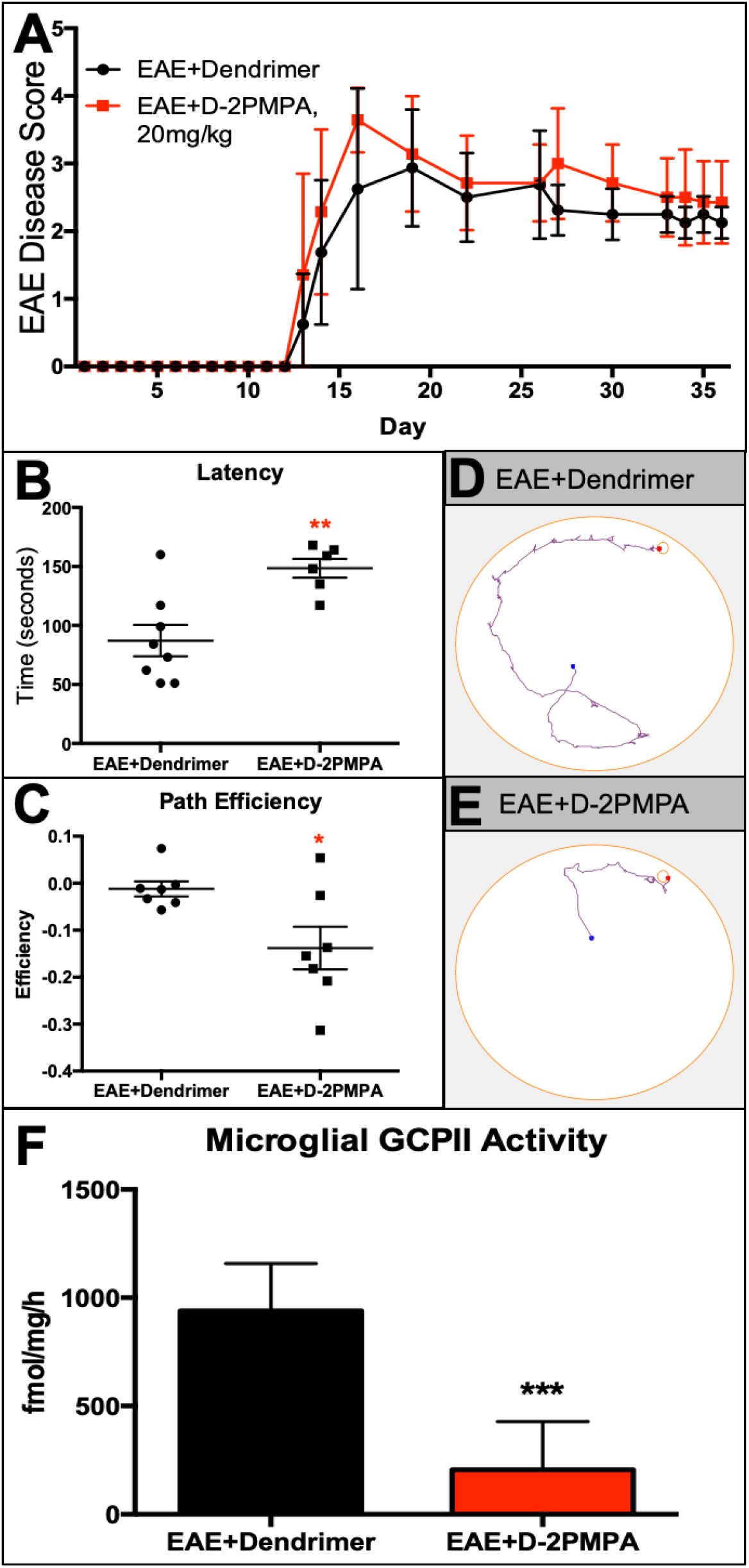
D-2PMPA does not affect physical severity of EAE, but selectively improves cognitive function and inhibits glial GCPII activity. **A.** Biweekly 20mg/kg D-2PMPA treatment initiated at the onset of physical disease had no impact on EAE physical disease severity. At 5 weeks post-immunization, EAE mice were tested in the Barnes maze. D-2PMPA caused a significant improvement in cognition as measured by the change between the first and last trial in **B.** latency to find the target and **C.** path efficiency. **D-E.** Representative track plot reports from the final trial in Barnes maze testing illustrate improved performance in mice treated with D-2PMPA versus vehicle dendrimer. Experiments were performed in duplicate with n= 7-8 mice, and the data are plotted as mean ± SEM of a representative experiment. **F.** Following Barnes maze testing, EAE mice received a terminal dose of dendrimer or D-2PMPA and were sacrificed 24 hours later. CD11b+ cells isolated from hippocampi showed a significant 4.5-fold decrease of GCPII activity. Significantly different from EAE + dendrimer at P<0.001(***).

Following 3 weeks of biweekly vehicle or D-2PMPA treatment, cognitive function was evaluated using the Barnes maze test in the same cohorts of mice monitored for disease score. Mice treated with D-2PMPA demonstrated superior learning and memory as evidenced by significantly elevated primary latency delta (first trial latency – final trial latency, **Figure 7B**, P<0.01). Path efficiency is the distance from the center starting point of the maze divided by the total distance traveled, with a score approaching 1 indicating the most efficient path to the target. The path efficiency delta (first trial efficiency – final trial efficiency, **Figure 7C**, P<0.05) was significantly lower in D-2PMPA mice, demonstrating more proficient learning versus EAE+dendrimer mice. Paths in the maze were recorded and automatically tracked, and representative track plots from the final trial of Barnes maze testing show superior performance due to D-2PMPA treatment (**Figure 7D, E**).

#### 3.4.03 D-2PMPA inhibits GCPII activity in hippocampal CD11b+ cells from EAE mice

To confirm target engagement following D-2PMPA treatment, GCPII enzymatic activity levels were measured in microglial-enriched cells of animals that completed Barnes maze cognitive testing. D-2PMPA treatment led to a near complete inhibition of enzymatic activity, with a >75% reduction versus vehicle dendrimer (**Figure 7F**, P<0.001).

## 4. Discussion

We synthesized a GCPII inhibitor dendrimer-drug conjugate by attaching 2-PMPA to the surface of hydroxyl PAMAM dendrimers, and evaluated its ability to deliver 2-PMPA to activated glia and improve cognition in the EAE model of MS. The structure of 2-PMPA includes a dicarboxylic acid and a polar phosphonate making it highly hydrophilic, resulting in low membrane permeability, negligible oral bioavailability, and very limited brain penetration^28^. In preclinical models of neurological diseases where inhibition of GCPII has shown therapeutic potential, 2-PMPA is efficacious only after very high systemic doses (i.p.) or direct brain injection, and is therefore not optimal for chronic dosing in patients^8,9^. To circumvent these issues and enhance brain uptake the drug was covalently conjugated to the surface of hydroxyl PAMAM dendrimers using a highly efficient, robust and atom economical CuAAC approach^29,30^. We show that the conjugation of 2-PMPA to dendrimers facilitates the targeted delivery of 2-PMPA to activated glial cells from systemic administration, and that 2-PMPA delivery to glia via dendrimers allows for only twice weekly administration of a 20mg/kg dose to significantly improve cognitive impairment in EAE, accounting for a 17.5-fold reduction in the free 2-PMPA required to achieve similar behavioral outcomes^8^. This suggests that targeted delivery with the dendrimer reduces the systemic drug dose required, potentially improving the side effect profile and the opportunity for translation.

The present study is the first to report that dendrimers are a potent means of delivering drugs for the treatment of MS-related cognitive impairment. Synthesis of fluorescently labeled D-2PMPA allowed for visualization of dendrimer uptake in both cultured glial cells and in EAE brains. Similar to previously published reports, we observed very limited uptake of dendrimer conjugates into astrocytes as compared with microglia in culture studies, and no uptake into astrocytes in EAE brains^23,31^. To our knowledge, only one other study has tested a dendrimer drug delivery system in the EAE mouse model of MS, with positive anti-inflammatory results resulting from treatment with an amino-bis(methylene phosphonate)-capped dendrimer^32^. Dendrimers have also proven efficacious in targeting glial drug delivery in other models of neurological disease, including inflammatory preterm birth injury^13^, cerebral palsy^14^, and Rett Syndrome^16^. Because glial cells are implicated in the pathogenesis of many neurological diseases and models, including EAE and MS^33^, dendrimer drug delivery has the potential to assist in the generation of new treatments for many diseases with scarce and insufficient treatment options. Furthermore, the clinical translation of drugs with either poor pharmacokinetic profiles that prevent CNS delivery, such as GCPII inhibitors, or toxic side effects at pharmacologically relevant doses can now be potentially revisited.

EAE is a highly inflammatory disease, and previous studies have shown that both TNFα and iNOS can contribute to EAE disease progression^34,35^. In the present study, we show that D-2PMPA downregulates both of these pro-inflammatory markers in glial cultures, along with an upregulation of anti-inflammatory markers, so we were surprised to find that no changes in EAE scores were observed due to D-2PMPA treatment. This finding, however, is in line with our previous observations that free 2-PMPA treatment had no effect on EAE disease score or inflammation^8,9^. Clearly, targeted delivery of the drug with the dendrimer has improved the anti-inflammatory activity of the drug. Others have reported an opposite effect^36^, and we acknowledge that it is possible that with a modified treatment paradigm (i.e. earlier or more frequent dosing, or combination therapy) EAE physical severity would be impacted. Future studies will explore this idea.

Although physical severity was unaltered, a significant improvement in cognition in EAE mice was detected due to D-2PMPA treatment. It is possible that these improvements are due to anti-inflammatory effects of D-2PMPA in the brain. In addition to contributing to EAE disease progression, elevations in TNFα cause synaptic instability and have been linked to cognitive dysfunction in EAE^37^. Additionally, reductions in TNFα and iNOS levels are both associated with improved cognition in rats^38^. It is also possible that a decrease in glutamate excitotoxicity and/or an upregulation of NAAG levels contributed to learning and memory improvements in D-2PMPA-treated EAE mice. GCPII is located on the surface of glial cells in the extrasynaptic space, with an extracellular active site^5^. Under conditions of high synaptic activity, such as neuropathological injury or disease, the release of NAAG and subsequent cleavage of the neuropeptide by GCPII is elevated, which in turn results in increased extrasynaptic glutamate levels that can lead to excitotoxicity. In line with this, administration of the GCPII inhibitor 2-PMPA reverses ischemia-induced elevations in glutamate and subsequent neurotoxicity^39^. Previous studies in our laboratory have shown that TGFβ, found here to be upregulated up to 16-fold following *in vitro* D-2PMPA treatment, is required for GCPII to afford neuroprotection^40^. NAAG is an agonist at mGluR3, and activation of mGluR3, found here to be upregulated by D-2PMPA, both enhances the release of neurotrophic factors^41^ and decreases excitatory neurotransmitter release via negative feedback to presynaptic neurons28,42. This mechanism has been confirmed by multiple laboratories that have shown that the positive effects of GCPII inhibition can be reversed following administration of mGluR3 antagonists^7,28,43^. In agreement, independent laboratories have shown similar positive relationships between brain NAAG levels and cognitive function in other neurological diseases, including schizophrenia^44^, and reductions in brain NAAG levels where cognitive impairment is a hallmark attribute, such as Alzheimer’s and Huntington’s diseases^45,46^. It is likely that a combination of anti-inflammatory properties, decreased glutamate levels, and increased NAAG levels contribute to the procognitive effects of D-2PMPA in EAE, and future studies will further explore these mechanistic possibilities.

As treatments that target MS-related disability improve and lifespan is prolonged, it is reasonable to predict that comorbidities such as cognitive impairment will manifest to a greater degree. In turn, the negative effects of cognitive impairment on social well-being and financial productivity are therefore likely to magnify. While small to moderate effects have been observed due to computerized cognitive training^47^, no drug therapies are currently designed and approved to treat cognitive impairment in MS. Inhibition of GCPII via targeted dendrimer drug delivery, therefore, has the potential to serve as the first therapeutic strategy to target MS-related cognitive impairment. The improved cognitive effects with D-2PMPA were achieved at a 17.5-fold lower drug levels, suggesting that the side effect profile will be significantly improved for the conjugate (cleared intact through the kidney), providing a larger therapeutic window for potential translation to humans.

## 5. Conclusions

Using hydroxyl PAMAM dendrimers, we have developed a potent brain penetrating GCPII inhibitor D-2PMPA, which enables the targeted delivery of a GCPII inhibitor to activated glial cells. We show that systemic D-2PMPA inhibits >75% of the GCPII enzymatic activity in CD11b+ cells in EAE brains and improved the cognition function in EAE mice at a 17.5-fold lower dose compared to the free drug. Taken together, these encouraging results suggest that dendrimer-targeted GCPII inhibition could be a potent therapeutic strategy for the improvement of cognition in MS patients.

## Data Availability

Data will be made available upon request.

## References

1. Benedict RH, Wahlig E, Bakshi R, et al. Predicting quality of life in multiple sclerosis: accounting for physical disability, fatigue, cognition, mood disorder, personality, and behavior change. J Neurol Sci. 2005;231(1-2):29–34.

2. Kobelt G, Thompson A, Berg J, et al. New insights into the burden and costs of multiple sclerosis in Europe. Mult Scler. 2017;23(8):1123–1136.

3. Campbell J, Rashid W, Cercignani M, Langdon D. Cognitive impairment among patients with multiple sclerosis: associations with employment and quality of life. Postgrad Med J. 2017;93(1097):143–147.

4. van Gorp DAM, van der Hiele K, Heerings MAP, et al. Cognitive functioning as a predictor of employment status in relapsing-remitting multiple sclerosis: a 2-year longitudinal study. Neurol Sci. 2019.

5. Vornov JJ, Hollinger KR, Jackson PF, et al. Still NAAG’ing After All These Years: The Continuing Pursuit of GCPII Inhibitors. Adv Pharmacol. 2016;76:215–255.

6. Neale JH, Bzdega T, Wroblewska B. N-Acetylaspartylglutamate: the most abundant peptide neurotransmitter in the mammalian central nervous system. J Neurochem. 2000;75(2):443–452.

7. Olszewski RT, Bzdega T, Neale JH. mGluR3 and not mGluR2 receptors mediate the efficacy of NAAG peptidase inhibitor in validated model of schizophrenia. Schizophr Res. 2012;136(1-3):160–161.

8. Rahn KA, Watkins CC, Alt J, et al. Inhibition of glutamate carboxypeptidase II (GCPII) activity as a treatment for cognitive impairment in multiple sclerosis. Proc Natl Acad Sci U S A. 2012;109(49):20101–20106.

9. Hollinger KR, Alt J, Riehm AM, Slusher BS, Kaplin AI. Dose-dependent inhibition of GCPII to prevent and treat cognitive impairment in the EAE model of multiple sclerosis. Brain Res. 2016;1635:105–112.

10. Majumder J, Taratula O, Minko T. Nanocarrier-based systems for targeted and site specific therapeutic delivery. Adv Drug Deliv Rev. 2019;144:57–77.

11. Anselmo AC, Mitragotri S. Nanoparticles in the clinic. Bioeng Transl Med. 2016;1(1):10–29.

12. Leiro V, Santos SD, Lopes CDF, Pego AP. Dendrimers as Powerful Building Blocks in Central Nervous System Disease: Headed for Successful Nanomedicine. Advanced Functional Materials. 2017;28(12).

13. Lei J, Rosenzweig JM, Mishra MK, et al. Maternal dendrimer-based therapy for inflammation-induced preterm birth and perinatal brain injury. Sci Rep. 2017;7(1):6106.

14. Kannan S, Dai H, Navath RS, et al. Dendrimer-based postnatal therapy for neuroinflammation and cerebral palsy in a rabbit model. Sci Transl Med. 2012;4(130):130ra146.

15. Wang B, Navath RS, Romero R, Kannan S, Kannan R. Anti-inflammatory and anti-oxidant activity of anionic dendrimer-N-acetyl cysteine conjugates in activated microglial cells. Int J Pharm. 2009;377(1-2):159–168.

16. Nance E, Kambhampati SP, Smith ES, et al. Dendrimer-mediated delivery of N-acetyl cysteine to microglia in a mouse model of Rett syndrome. J Neuroinflammation. 2017;14(1):252.

17. Nino DF, Zhou Q, Yamaguchi Y, et al. Cognitive impairments induced by necrotizing enterocolitis can be prevented by inhibiting microglial activation in mouse brain. Sci Transl Med. 2018;10(471).

18. Sharma A, Liaw K, Sharma R, Zhang Z, Kannan S, Kannan RM. Targeting Mitochondrial Dysfunction and Oxidative Stress in Activated Microglia using Dendrimer-Based Therapeutics. Theranostics. 2018;8(20):5529–5547.

19. Sharma R, Kim SY, Sharma A, et al. Activated Microglia Targeting Dendrimer-Minocycline Conjugate as Therapeutics for Neuroinflammation. Bioconjug Chem. 2017;28(11):2874–2886.

20. Sharma R, Sharma A, Kambhampati SP, et al. Scalable synthesis and validation of PAMAM dendrimer-N-acetyl cysteine conjugate for potential translation. Bioeng Transl Med. 2018;3(2):87–101.

21. Zhang Z, Bassam B, Thomas AG, et al. Maternal inflammation leads to impaired glutamate homeostasis and up-regulation of glutamate carboxypeptidase II in activated microglia in the fetal/newborn rabbit brain. Neurobiol Dis. 2016;94:116–128.

22. Sharma A, Porterfield JE, Smith E, Sharma R, Kannan S, Kannan RM. Effect of mannose targeting of hydroxyl PAMAM dendrimers on cellular and organ biodistribution in a neonatal brain injury model. J Control Release. 2018;283:175–189.

23. Alnasser Y, Kambhampati SP, Nance E, et al. Preferential and Increased Uptake of Hydroxyl-Terminated PAMAM Dendrimers by Activated Microglia in Rabbit Brain Mixed Glial Culture. Molecules. 2018;23(5).

24. Paxinos G, Franklin KBJ. Paxinos and Franklin’s the mouse brain in stereotaxic coordinates. 4th ed. Amsterdam: Elsevier/Academic Press; 2013.

25. Zhu X, Nedelcovych MT, Thomas AG, et al. JHU-083 selectively blocks glutaminase activity in brain CD11b(+) cells and prevents depression-associated behaviors induced by chronic social defeat stress. Neuropsychopharmacology. 2019;44(4):683–694.

26. Robinson MB, Blakely RD, Couto R, Coyle JT. Hydrolysis of the brain dipeptide N-acetyl-L-aspartyl-L-glutamate. Identification and characterization of a novel N-acetylated alpha-linked acidic dipeptidase activity from rat brain. J Biol Chem. 1987;262(30):14498–14506.

27. Rojas C, Frazier ST, Flanary J, Slusher BS. Kinetics and inhibition of glutamate carboxypeptidase II using a microplate assay. Anal Biochem. 2002;310(1):50–54.

28. Adedoyin MO, Vicini S, Neale JH. Endogenous N-acetylaspartylglutamate (NAAG) inhibits synaptic plasticity/transmission in the amygdala in a mouse inflammatory pain model. Mol Pain. 2010;6:60.

29. Kolb HC, Finn MG, Sharpless KB. Click Chemistry: Diverse Chemical Function from a Few Good Reactions. Angew Chem Int Ed Engl. 2001;40(11):2004–2021.

30. Thirumurugan P, Matosiuk D, Jozwiak K. Click chemistry for drug development and diverse chemical-biology applications. Chem Rev. 2013;113(7):4905–4979.

31. Nemeth CL, Drummond GT, Mishra MK, et al. Uptake of dendrimer-drug by different cell types in the hippocampus after hypoxic-ischemic insult in neonatal mice: Effects of injury, microglial activation and hypothermia. Nanomedicine. 2017;13(7):2359–2369.

32. Hayder M, Varilh M, Turrin CO, et al. Phosphorus-Based Dendrimer ABP Treats Neuroinflammation by Promoting IL-10-Producing CD4(+) T Cells. Biomacromolecules. 2015;16(11):3425–3433.

33. Duffy SS, Lees JG, Moalem-Taylor G. The contribution of immune and glial cell types in experimental autoimmune encephalomyelitis and multiple sclerosis. Mult Scler Int. 2014;2014:285245.

34. Valentin-Torres A, Savarin C, Hinton DR, Phares TW, Bergmann CC, Stohlman SA. Sustained TNF production by central nervous system infiltrating macrophages promotes progressive autoimmune encephalomyelitis. J Neuroinflammation. 2016;13:46.

35. Ding M, Zhang M, Wong JL, Rogers NE, Ignarro LJ, Voskuhl RR. Antisense knockdown of inducible nitric oxide synthase inhibits induction of experimental autoimmune encephalomyelitis in SJL/J mice. J Immunol. 1998;160(6):2560–2564.

36. Ha D, Bing SJ, Ahn G, et al. Blocking glutamate carboxypeptidase II inhibits glutamate excitotoxicity and regulates immune responses in experimental autoimmune encephalomyelitis. FEBS J. 2016;283(18):3438–3456.

37. Yang G, Parkhurst CN, Hayes S, Gan WB. Peripheral elevation of TNF-alpha leads to early synaptic abnormalities in the mouse somatosensory cortex in experimental autoimmune encephalomyelitis. Proc Natl Acad Sci U S A. 2013;110(25):10306–10311.

38. Sun J, Zhang S, Zhang X, Zhang X, Dong H, Qian Y. IL-17A is implicated in lipopolysaccharide-induced neuroinflammation and cognitive impairment in aged rats via microglial activation. J Neuroinflammation. 2015;12:165.

39. Slusher BS, Vornov JJ, Thomas AG, et al. Selective inhibition of NAALADase, which converts NAAG to glutamate, reduces ischemic brain injury. Nat Med. 1999;5(12):1396–1402.

40. Thomas AG, Liu W, Olkowski JL, et al. Neuroprotection mediated by glutamate carboxypeptidase II (NAALADase) inhibition requires TGF-beta. Eur J Pharmacol. 2001;430(1):33–40.

41. Thomas AG, Olkowski JL, Slusher BS. Neuroprotection afforded by NAAG and NAALADase inhibition requires glial cells and metabotropic glutamate receptor activation. Eur J Pharmacol. 2001;426(1-2):35–38.

42. Wroblewska B, Wegorzewska IN, Bzdega T, Olszewski RT, Neale JH. Differential negative coupling of type 3 metabotropic glutamate receptor to cyclic GMP levels in neurons and astrocytes. J Neurochem. 2006;96(4):1071–1077.

43. Zhong C, Zhao X, Van KC, et al. NAAG peptidase inhibitor increases dialysate NAAG and reduces glutamate, aspartate and GABA levels in the dorsal hippocampus following fluid percussion injury in the rat. J Neurochem. 2006;97(4):1015–1025.

44. Jessen F, Fingerhut N, Sprinkart AM, et al. N-acetylaspartylglutamate (NAAG) and N-acetylaspartate (NAA) in patients with schizophrenia. Schizophr Bull. 2013;39(1):197–205.

45. Jaarsma D, Veenma-van der Duin L, Korf J. N-acetylaspartate and N-acetylaspartylglutamate levels in Alzheimer’s disease post-mortem brain tissue. J Neurol Sci. 1994;127(2):230–233.

46. Passani LA, Vonsattel JP, Carter RE, Coyle JT. N-acetylaspartylglutamate, N-acetylaspartate, and N-acetylated alpha-linked acidic dipeptidase in human brain and their alterations in Huntington and Alzheimer’s diseases. Mol Chem Neuropathol. 1997;31(2):97–118.

47. Lampit A, Heine J, Finke C, et al. Computerized Cognitive Training in Multiple Sclerosis: A Systematic Review and Meta-analysis. Neurorehabil Neural Repair. 2019:1545968319860490.

